# Inconsistent effects of urbanization on amphibian genetic diversity

**DOI:** 10.1101/2020.08.16.253104

**Authors:** Chloé Schmidt, Colin J Garroway

## Abstract

Habitat loss and fragmentation are leading causes of vertebrate population declines. These declines are thought to be partly due to decreased connectivity and habitat loss reducing population sizes in human transformed habitats. With time this can lead to reduced effective population size and genetic diversity which restricts the ability of wildlife to cope with environmental change through genetic adaptation. However, it is not well understood whether these effects are generally applicable across taxa. Here, we repurposed and synthesized raw microsatellite data from online repositories from 19 amphibian species sampled at 554 sites in North America. For each site, we estimated gene diversity, allelic richness, effective population size, and population differentiation. Using binary urban-rural census designations, and continuous measures of human population density and the Human Footprint Index, we tested for generalizable effects of human land use on amphibian genetic diversity. We found no consistent relationships for any of our genetic metrics. While we did not detect directional effects for most species, a few generalist species responded positively to urbanization. These results contrast with consistent negative effects of urbanization in mammals and species specific positive and negative effects in birds. In the context of widespread amphibian declines, our results suggest that habitat loss in human transformed habitats is a more immediate concern than declining genetic diversity in populations that persist.

## Introduction

Populations of terrestrial vertebrates are experiencing declines globally due to habitat loss and the conversion of natural land for human purposes (WWF, 2018). Proportionately more amphibians (41%) are threatened with extinction than are mammals (25%), reptiles (22%), or birds (13%) (Hoffmann et al., 2010). Due to data deficiencies for many amphibians this risk is likely underestimated (Stuart et al., 2004). Habitat transformation by humans is among the foremost threats to vertebrate biodiversity, in terms of both species losses and declining population sizes (Hamer & McDonnell, 2008). Amphibians are clearly sensitive to habitat degradation (Collins, Crump, & Lovejoy III, 2009; Hamer & McDonnell, 2008), but we do not yet know whether amphibians exhibit generalizable population genetic responses to urbanization and other human land uses.

Genetic diversity is the most fundamental level of biodiversity. It is important for conservation because it reflects a population’s ability to adaptively respond to environmental change (Pereira et al., 2013). Small populations typically have lower genetic diversity, higher rates of inbreeding depression, experience stronger effects of genetic drift, and have a reduced capacity to purge deleterious alleles and respond to selection pressures — this reduces long-term population viability (Frankham, 1995). These genetic effects are a consequence of effective population size (Falconer & Mackay, 1996). The effective population size is the size of an ideal population with constant size, randomly mating individuals, and non-overlapping generations that produces the same rate of genetic drift as the measured population. Over time, the random loss or fixation of alleles caused by drift becomes the dominant force driving evolutionary change in populations with small effective sizes, regardless of census population size.

Habitat fragmentation and loss due to urbanization is a major cause of population decline (WWF, 2018). This is because it divides populations and impedes dispersal between them, leading to greater genetic differentiation and smaller effective population sizes, which in turn reduces genetic diversity and strengthens genetic drift (Johnson & Munshi-South, 2017; Miles, Rivkin, Johnson, Munshi-South, & Verrelli, 2019; Schmidt, Domaratzki, Kinnunen, Bowman, & Garroway, 2020). Mammals are the most well-studied taxa in urban evolutionary ecology (Miles et al., 2019; Schell, 2018) and they tend to adhere to the expectation of increased drift and genetic differentiation in disturbed environments (DiBattista, 2008; Schmidt et al., 2020). Whether these effects are generalizable across taxa is unclear (Miles et al., 2019). Birds, for example, did not show consistent effects of urbanization among species in North America (Schmidt et al., 2020)—but increased vagility due to the ability to fly seems likely to explain much of this difference. In amphibians, species-specific studies have tended to find lower genetic diversity and higher population differentiation for populations in urban areas (Arens et al., 2007; Miraldo et al., 2016; Munshi-South, Zak, & Pehek, 2013; Noёl, Ouellet, Galois, & Lapointe, 2007). Amphibians’ mode of locomotion and sensitivity to environmental conditions suggests that species, in general, might respond to the effects of urban habitat fragmentation negatively, similar to mammals.

Urbanization has particularly negative consequences for amphibian populations due to their complex habitat requirements. Declines in species richness and abundance are associated with the reduction and isolation of wetlands, wetland vegetation, and forest cover (Collins et al., 2009; Hamer & McDonnell, 2008). Some species, especially those that rely on vernal pools, are unable to complete their life cycles in urban environments where aquatic and terrestrial habitats are separated by roads and other urban infrastructure (Hamer & McDonnell, 2008). Temporary wetlands are typically not a priority for habitat conservation, and due to their small size, can dry out easily and are more vulnerable to pollution in human-dominated habitats (Hamer & McDonnell, 2008). Locally, amphibians have patchy distributions that form metapopulation structures at regional scales. Limited population connectivity due to wetland destruction and fragmentation can sever connections within the metapopulation network, causing population isolation. The genetic diversity of amphibian populations could thus be especially vulnerable to continued land-use change.

We tested for general effects of urbanization on gene diversity, allelic richness, effective population size, and population-specific F_ST_ across a sample of North American amphibians by repurposing archived georeferenced microsatellite data. We harvested data amounting to 13,680 individual genotypes from 19 species sampled at 554 sites in Canada and the United States (Table S1). Synthesizing and repurposing data collected for different questions is powerful because it is unlikely that study system selection, for instance a focus on systems where strong effects are suspected, would bias our findings. Additionally, this approach allowed us to consistently calculate both genetic and environmental variables that were not necessarily presented in the original papers. We focused on North America to control for effects of regional history on genetic diversity and population structure (Hewitt, 2000; Schmidt et al., 2020). Additionally, the southeastern United States is an important hotspot for salamander species diversity. We used three measures of landscape change that encompass different aspects of urbanization and human presence. Census-based urban/rural designation is a broad scale categorical measure of urbanization. We used human population density measured per square kilometer as a finer scale continuous measure of human presence. Finally, we used the Human Footprint Index, a more comprehensive fine-scale measure of human presence and disturbance which incorporates population density, built-up areas, night-time lights, land cover, and human access to coastlines, roads, railways and navigable rivers (WCS & CIESIN, 2005).

## Methods

### Data assembly

We conducted a systematic search of online repositories for raw microsatellite data in February 2019 using the DataONE interface (Jones et al., 2017) through R (R Core Team, 2013). We focused on microsatellites rather than single-nucleotide polymorphisms (SNPs) because they are widely used for genetic monitoring of wildlife populations, and raw data is commonly archived in online repositories (Miles et al., 2019). Neutral microsatellite markers provide good estimates of genome-wide diversity (Mittell, Nakagawa, & Hadfield, 2015) in addition to useful information about population demography. We searched for species names using a list of amphibian species native to North America from the IUCN Red List database (for example, “*Ambystoma maculatum*”), in addition to the terms: “microsat*”, “short tandem*”, or “single tandem*”. We obtained 51 search results. Twenty-three results were duplicates (owing to different data versions archived in the Dryad repository) and were removed (see Data S1 for complete list of search results).

We screened the remaining data for suitability according to the following criteria: raw, neutral microsatellite data available; species present in data were on search list; the location of data collection was North America; and that the original study design would not have effects on genetic diversity, such as recently bottlenecked or island populations. In some cases multiple datasets from one research group included the same species sampled at overlapping sites. In these cases we kept the dataset with a greater number of sample locations to avoid resampling the same populations. Ultimately 20 out of 28 unique datasets were retained for reanalysis.

Next, we imported selected datasets into R using the adegenet package (Jombart et al., 2017). We calculated two measures of genetic diversity, gene diversity (Nei, 1973) and allelic richness, effective population size, and population-specific F_ST_ (Weir & Goudet, 2017). Gene diversity is a measure of the spread and evenness of alleles in a population. It primarily takes into account allele frequencies and is therefore is only minimally affected by different sample sizes (Charlesworth & Charlesworth, 2010). Allelic richness is a count of alleles in a population that is typically standardized to a minimum sample size by rarefaction (Leberg, 2002). Here, we set a minimum sample size of 5 individuals. We estimated the effective population size of the parental generation using the linkage disequilibrium method (Waples & Do, 2010) in Neestimator v. 2 (Do et al., 2014; Waples & Do, 2008). This method is among the more accurate, especially for smaller effective population sizes (Waples & Do, 2010). However, in large populations signals of drift are overwhelmed by sampling error which hinders estimation of effective population size. If too few individuals or loci were sampled to provide information on the strength of genetic drift, an estimate of infinity is returned. We excluded these estimates from our analyses. Finally, population-specific F_ST_ is a measure of population differentiation which estimates how far populations in a sample have diverged from a common ancestor (Weir & Goudet, 2017). This method requires at least 2 populations per sample, and we were unable to estimate F_ST_ when this requirement was not met.

### Human influence

We measured the effects of humans and urbanization in three ways. First, we assigned each site to urban or non-urban categories depending on whether they overlapped with census-designated urban areas and population centers in the United States and Canada (Statistics Canada, 2016; U.S. Census Bureau, 2016). Next, we measured the average human population density per km^2^ and Human Footprint Index (WCS & CIESIN, 2005) at each site within a buffer zone. We used 1, 5, 10, and 15 km buffers around each site to test whether relationships between genetic metrics and human presence were scale-dependent. Responses to population density varied consistently in strength and direction regardless of scale, and effects of the human footprint began to be distinguishable with a 10 km buffer (Fig. S1). We report results from the 10 km buffer in the main text as this allowed for comparison to results for mammals and birds from previous work using this analytical approach (Schmidt et al., 2020). Full results are presented in SI Fig. S1.

### Analysis

We examined the effects of human land transformation on genetic diversity (gene diversity and allelic richness), effective population size, and population differentiation (population-specific F_ST_). However, there are likely spatial patterns in the distribution of genetic diversity present in the data by virtue of site placement (closer sites are more similar), population structure and demography, or historical factors such as climate and glaciation cycles, or species life history (e.g. dispersal distance). Following Schmidt et al. (2020), we controlled for factors that could create spatial structure in the data with distance-based Moran’s eigenvector maps (MEMs). This method produces spatial eigenvectors with positive eigenvalues which are directly proportional to Moran’s *I* index of spatial correlation. MEMs capture spatial variation across all scales present in the data, and can be used to control for spatial patterns caused by unknown factors (Manel, Poncet, Legendre, Gugerli, & Holderegger, 2010). We computed MEMs in the adespatial package (version 0.3.8) (Dray et al., 2017) and used a forward selection procedure to select MEMs which described important spatial variation in gene diversity, allelic richness, effective population size, and population-specific F_ST_ (Blanchet, Legendre, & Borcard, 2008).

Next, we tested for effects of human presence on population genetic composition using hierarchical models in a Bayesian framework. For each measure of genetic composition (gene diversity, effective population size, and population-specific F_ST_) we fitted a series of four linear mixed models. Three models each included a measure of human presence (urban/rural classification, human population density, and Human Footprint Index) and selected MEMs, while the fourth model was a null model which included MEMs only or were intercept-only models if no spatial autocorrelation was detected. We accounted for species differences by including species as a random effect allowing slopes and intercepts to vary. Random slope and intercept models can be interpreted as first estimating the effects of human presence across populations within species, then generalizing coefficient estimates across species. Effective population size and human population density were log-transformed, and all variables were scaled and centered before analysis allowing us to compare effect sizes across models. Bayesian regression models were run using brms (Bürkner, 2019) with 4 chains, 5000 iterations (1000 burn-in iterations), and default uniform priors. We calculated marginal (R^2^_m_) and conditional R^2^ (R^2^_c_) values to determine the proportion of variation explained by fixed effects, and fixed and random effects together, respectively.

## Results

### Data synthesis

From the 20 previously published datasets, we obtained raw microsatellite genotypes from 13,680 individuals of 19 species sampled at 554 sites in Canada and the United States (Fig 1; Table S1). The median number of individuals sampled per site was 21 (range: 5 – 299), and the mean number of sampled loci was 11.15 (range: 5 – 20). The average gene diversity across species was 0.68 ± 0.16 SD, and for allelic richness was 5.16 ± 2.11 SD. Species-specific summaries are presented in Table S1.

**Figure 1.**
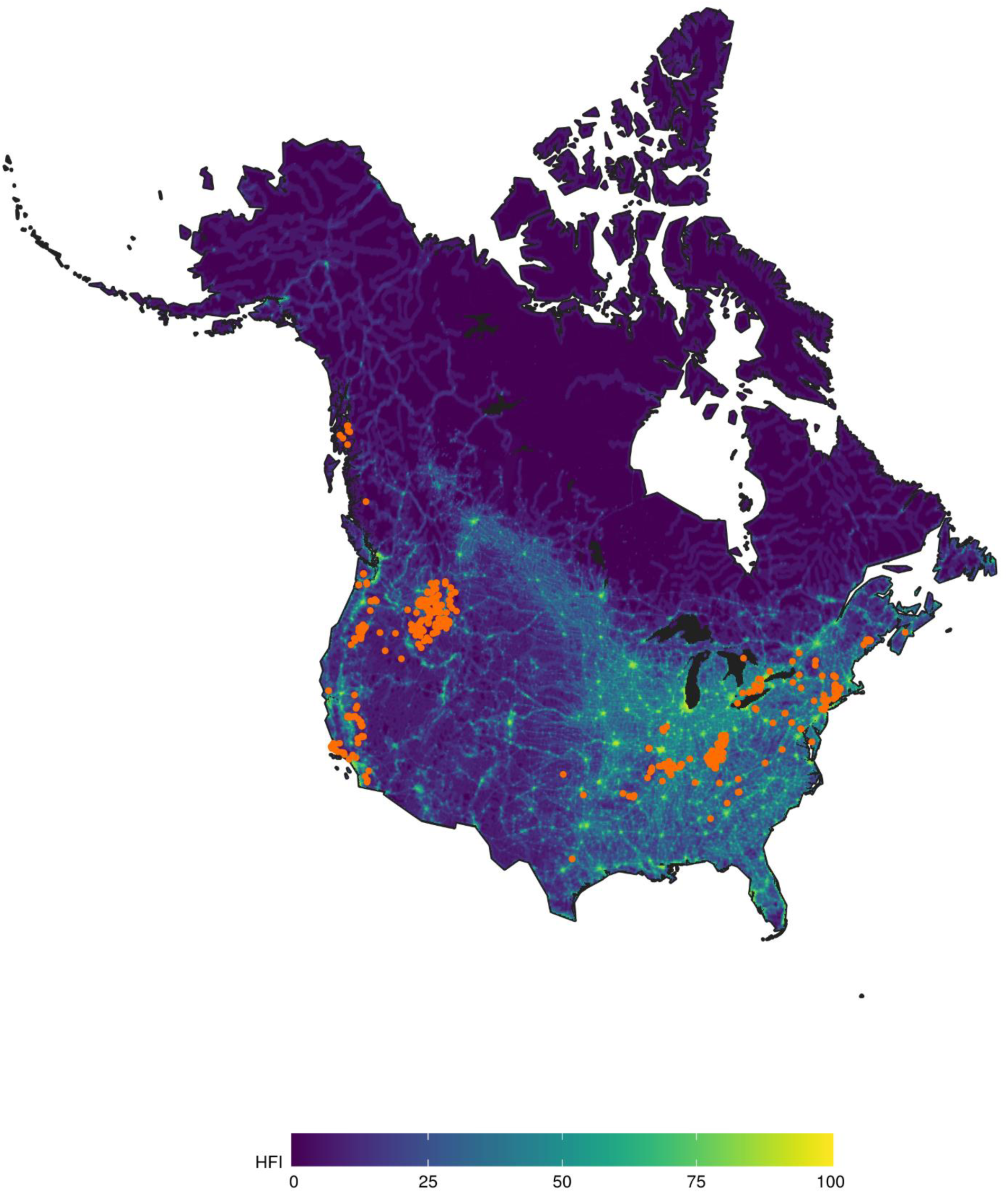
Map showing the locations of 554 sample sites from 19 species included in our analyses. We harvested raw microsatellite data sampled from each site to estimate gene diversity, allelic richness, effective population size, and population-specific F_ST_. Sites are overlaid on a map of the Human Footprint Index (HFI), which ranges from 0 (most wild) to 100 (most transformed).

We were able to measure population-specific F_ST_ for all but 2 sample sites where only a single site was sampled. Mean F_ST_ was 0.11 ± 0.15 SD. Finally, we were only able to estimate effective population size for 387 sites. The average effective population size was 188.27 ± 648.59 SD individuals, however, it varied widely between sites (range: 1.3 – 7847.8).

### Effects of urbanization

Human presence and urbanization did not tend to have strong effects on amphibian genetic diversity or population structure (Fig. 2, Table 1). Census-based urban/rural classification had no effects on any genetic metric. There was a trend of higher genetic diversity and effective population sizes, and lower population differentiation at sites with higher Human Footprint Index and population density. However, the only clear effects were a negative effect of Human Footprint Index on population-specific F_ST_ and a positive effect of human population density on effective population size (Fig. 2). Urban predictors had particularly low explanatory power for effective population size and allelic richness relative to null models (Table 1). For gene diversity, models with population density or Human Footprint Index—continuous measures of urbanization—explained the largest amount of variation (19% and 17% respectively; Table 1).

**Figure 2.**
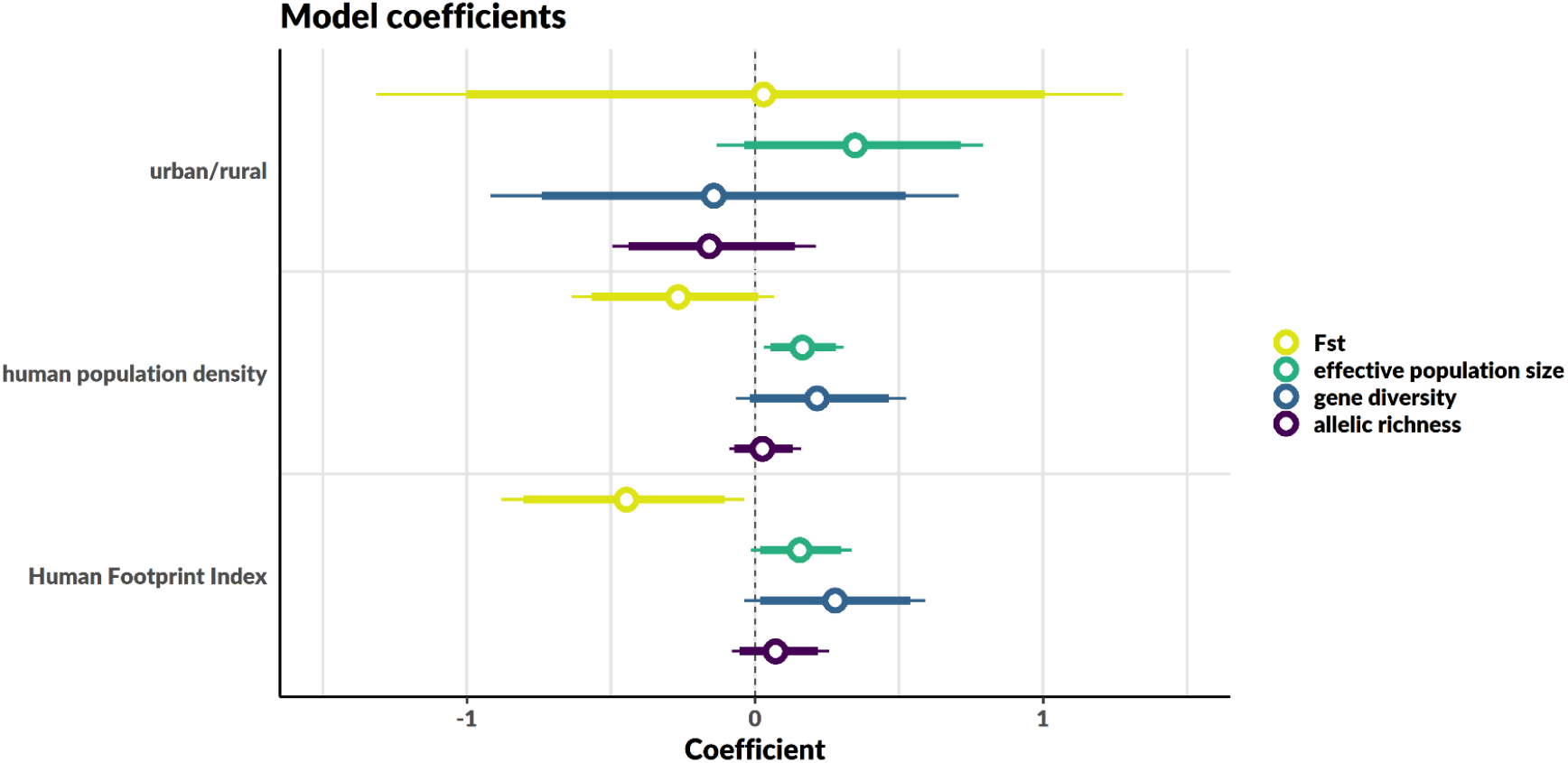
Coefficients from Bayesian linear mixed models. Effect sizes (open circles) are shown with 90% (bold lines) and 95% (thin lines) credible intervals. Note sample sizes differed between genetic measures (see Table 1). Effects appear generally small and inconsistent, however, there is a clear positive relationship between effective population size and human population density, and a clear negative relationship between population-specific F_ST_ and the Human Footprint Index.

**Table 1.** Model summaries for the effects of urbanization and human presence on population genetic composition. Four models were run for each response variable, three of which included a measure of human presence. The fourth was a null model including only spatial predictors as fixed effects if spatial autocorrelation was detected, otherwise was an intercept-only model. Marginal R^2^ refers to the proportion of variation explained by fixed effects, and conditional R^2^ is the variation explained by both fixed and random effects.

| Variable | Covariates | Coefficient (95% CI) | Marginal; Conditional $R^2$ |
| --- | --- | --- | --- |
| effective population size |  |  |  |
| n = 387 | urban/rural | 0.35 (-0.13 – 0.79) | 0.01; 0.32 |
|  | human population density | 0.16 (0.03 – 0.31) | 0.03; 0.32 |
|  | Human Footprint Index | 0.15 (-0.01 – 0.34) | 0.02; 0.32 |
|  | none | – | 0.00; 0.31 |
| gene diversity |  |  |  |
| n = 554 | urban/rural | -0.14 (-0.92 – 0.71) | 0.10; 0.74 |
|  | human population density | 0.21 (-0.07 – 0.52) | 0.19; 0.76 |
|  | Human Footprint Index | 0.28 (-0.04 – 0.59) | 0.17; 0.77 |
|  | none | – | 0.11; 0.74 |
| allelic richness |  |  |  |
| n = 554 | urban/rural | -0.16 (-0.50 – 0.21) | 0.03; 0.59 |
|  | human population density | 0.02 (-0.09 – 0.16) | 0.03; 0.59 |
|  | Human Footprint Index | 0.07 (-0.08 – 0.26) | 0.04; 0.60 |
|  | none | – | 0.02; 0.59 |
| $F_{ST}$ | | | |
| n = 552 | urban/rural | 0.03 (-1.32 – 1.28) | 0.15; 0.58 |
|  | human population density | -0.27 (-0.64 – 0.07) | 0.17; 0.60 |
|  | Human Footprint Index | -0.45 (-0.88 – -0.04) | 0.19; 0.62 |
|  | none | – | 0.18; 0.56 |

Effects of human presence on genetic diversity varied between species (Fig. S2). Genetic diversity of Cope’s giant salamander (*Dicamptodon copei*), the Rocky Mountain tailed frog (*Ascaphus montanus*), and the spring peeper (*Pseudacris crucifer*) tended to increase in more urban habitats whereas genetic diversity decreased with urbanization for northern dusky salamanders (*Desmognathus fuscus*) and California red-legged frogs (*Rana draytonii*) (Fig. S2). There was no clear direction of effect for the remaining 14 species.

## Discussion

In general, genetic diversity and population structure in North American amphibians were not consistently related to the measures of urbanization and human disturbance we tested. Responses to human presence measured by the Human Footprint Index and population density trended in the same directions. In contrast to our expectations, amphibians showed a general tendency towards lower levels of population differentiation, higher effective population sizes, and greater genetic diversity in transformed environments. Species-specific responses to urbanization (Fig. S2) suggest that pronounced positive effects in a few species drove this pattern. Cope’s giant salamander, the Rocky mountain tailed frog, and the spring peeper all had increased genetic diversity in more urban habitats. Several factors could have contributed to this result. Among amphibians, widely distributed generalist species, pond breeders, and species with aquatic development are more tolerant of habitat disturbance (Hamer & McDonnell, 2008; Nowakowski, Frishkoff, Thompson, Smith, & Todd, 2018). Relative to other species in our dataset, Rocky Mountain tailed frogs and pond breeding spring peepers both have two of the largest ranges. Other species with large ranges included spotted salamanders (*Ambystoma maculatum*), streamside salamanders (*A. barbouri*), and wood frogs (*Lithobates sylvaticus*), which were unaffected by landscape measures of disturbance (Fig. S2). Other studies with data not included in our synthesis have also found that genetic diversity did not decrease with urbanization in wood frogs (Furman, Scheffers, Taylor, Davis, & Paszkowski, 2016), fire salamanders (Lourenço, Álvarez, Wang, & Velo-Antón, 2017), and northern two-lined salamanders (Fusco, Pehek, & Munshi-South, 2020). Abundance in a species of frog related to the spring peeper, the boreal chorus frog (*P. maculata*), had a positive relationship with urbanization, which may be attributed to its preference for open habitats, or potential breeding habitats near roads (Browne, Paszkowski, Foote, Moenting, & Boss, 2009).

Amphibian abundances at sites can vary across orders of magnitude from year to year (Collins et al., 2009), so it may be that for a majority of species genetic signals of urbanization are swamped by population fluctuations. Indeed, allelic richness—the number of alleles—is more sensitive to population bottlenecks than is gene diversity which measures the evenness and spread of alleles (Corunet & Luikart, 1996). Allelic richness data might thus be noisier than gene diversity for amphibians. Highly variable population sizes could then cause the discrepancy in model explanatory power we find between these two measures of genetic diversity: proximity to humans and spatial patterns explained substantially more of the variation in gene diversity (10-19%) than allelic richness (2-4%; Table 1).

These findings should not be taken as indicating that the loss of genetic diversity due to urbanization is not a concern for amphibians. While in our North American data set genetic diversity tended to be positively associated with urbanization, this general trend was driven by three species. There were also species that lost diversity with increasing urbanization (northern dusky salamander; California red-legged frog). Similarly, loss of genetic diversity has also been reported among urban amphibians (Arens et al., 2007; Munshi-South et al., 2013; Noël & Lapointe, 2010). We also note many of our sample sites were located in natural and moderately transformed landscapes (Fig. S3), and relatively few species were sampled in highly urbanized habitats. Human-made ponds and still waters in moderately transformed areas (e.g., agricultural land, parks, golf courses) may provide important breeding habitats for amphibians (Babbitt & Tanner, 2000; Barry, Pauley, & Maerz, 2008; Brand & Snodgrass, 2010; Dimauro & Hunter, 2002; Nowakowski et al., 2018; Saarikivi, Knopp, Granroth, & Merilä, 2013). The only species with populations consistently sampled in highly urbanized sites was the northern dusky salamander, which showed a marked, albeit non-significant, decline in genetic diversity with increasing human disturbance (Fig. S2) (Munshi-South et al., 2013).

It appears that, in contrast to mammals (Schmidt et al., 2020), amphibians do not generally lose diversity in response to urbanization. Indeed, for some species there may be, to a point, positive effects of urbanization on genetic diversity. For mammals, barriers to dispersal in human-transformed environments such as roads and buildings restrict movement (Tucker et al., 2018). However, amphibian movements are naturally more constrained due to strict physiological and environmental requirements (e.g., proximity to aquatic and terrestrial environments, presence of ephemeral ponds, forest cover, and soil moisture), which also contributes to patchy population distributions (Collins et al., 2009). It is possible that habitat fragmentation and loss limits dispersal and reduces the probability of recolonizing areas where species were previously extirpated (Guzy et al., 2012). This would explain losses at species and population levels without our detecting declines in genetic diversity. Our results suggest that with regards to urbanization, the outcomes of habitat loss and degradation—such as local extinction—are of more immediate concern than gradual long-term declines owing to reduced genetic diversity in this taxon.

While patterns of reduced genetic diversity and increased differentiation in mammals are quite consistent (DiBattista, 2008; Schmidt et al., 2020; Tucker et al., 2018), effects of urbanization within other vertebrate classes seem to depend more on the species in question. Bird species responded to urbanization, but responses were equally likely to be negative or positive (Schmidt et al., 2020). In birds, demographic changes in survival and reproduction in response to urbanization differ widely between species (Marzluff, 2001). Species’ life histories and ecological needs play an integral role in how species cope with land use change. In light of previous findings (DiBattista, 2008; Miles et al., 2019; Schmidt et al., 2020) and our results for amphibians, it appears that there is no one size fits all answer to questions about the effects of urbanization on genetic diversity (Miles et al., 2019). Although the direction of effect might vary, it is clear that urbanization alters the genetics of wildlife populations quite broadly.

## Acknowledgements

We thank Mitchell Green for his assistance collecting and formatting raw microsatellite data from online repositories. We would also like to thank the authors of the original studies whose data contributed to this work. C.S. and C.J.G. were supported by a Natural Sciences and Engineering Research Council of Canada Discovery Grant to C.J.G. C.S. was also supported by a U. Manitoba Graduate Fellowship, and a U. Manitoba Graduate Enhancement of Tri-council funding grant to C.J.G.

## Author contributions

CS and CJG conceived of the study. CS collected and processed genetic data. CS performed the analysis with input from CJG. CS wrote the first draft of the manuscript, and CJG contributed to editing subsequent drafts.

## Data accessibility

The raw data used in this study are available from public repositories, see Table S1 for a complete list of references. Synthesized data will be made available in the Dryad Digital Repository upon acceptance.

## Supplementary Information

**Contents:**

Figures S1–3

Table S1

**Figure S1.**
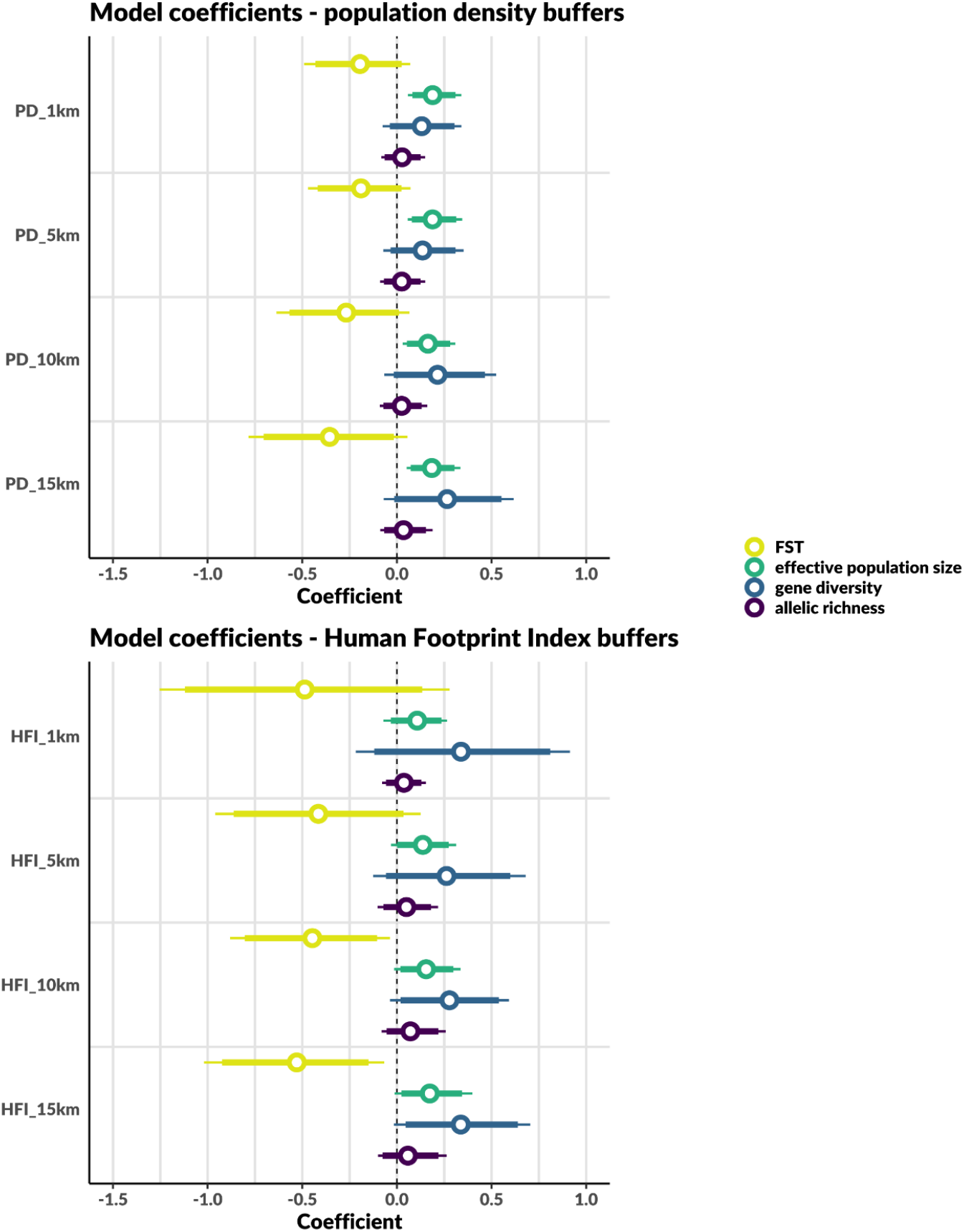
Effects of spatial scale. We extracted continuous measures of human presence within 1, 5, 10, and 15 km buffers around each site to test the effects of spatial scale on the relationships between genetic metrics and human presence. Responses to population density vary consistently in strength and direction regardless of scale, and effects of human footprint begin to be distinguishable with a 10 km buffer. We therefore selected a 10 km buffer to report in the main text for both variables.

**Figure S2.**
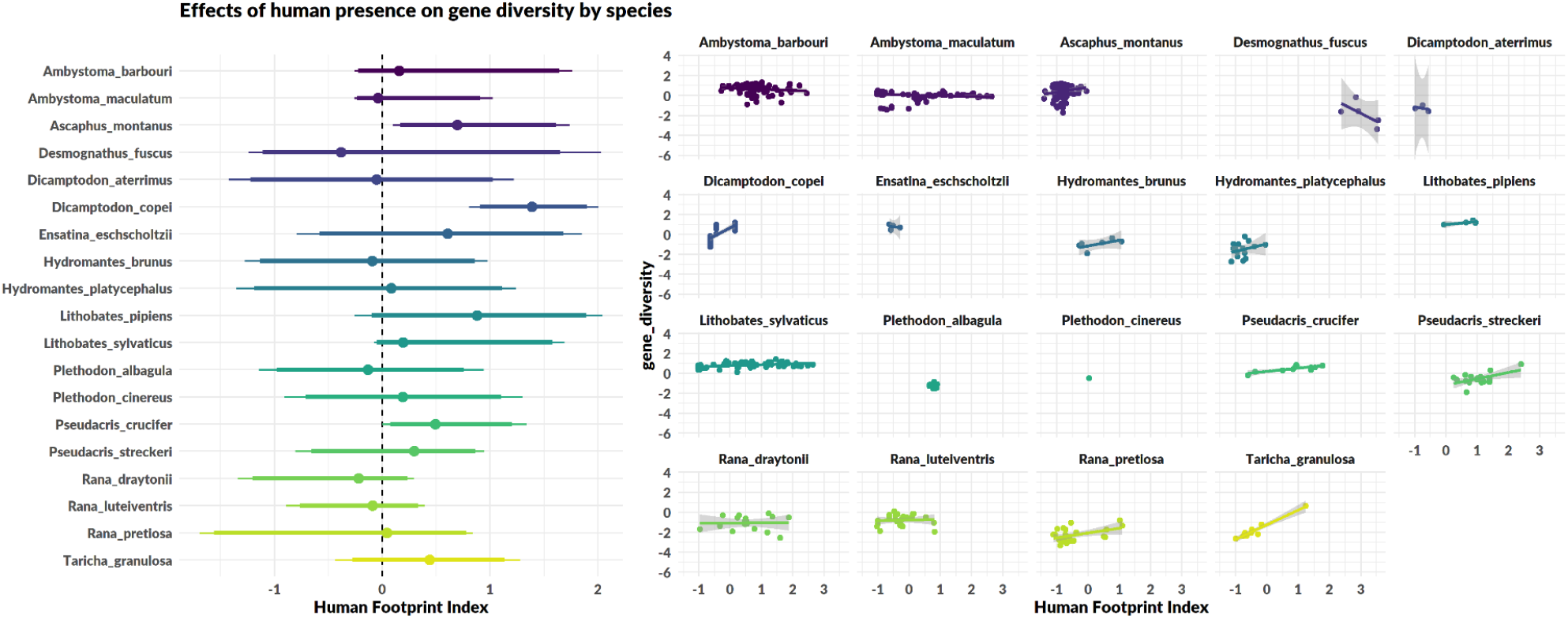
Species-specific effects of the Human Footprint Index on genetic diversity. Few species respond positively to urbanization (Rocky mountain tailed frog, *Ascaphus montanus*; Cope’s giant salamander, *Dicamptodon copei*; and spring peeper *Pseudacris crucifer*). In most species however, genetic diversity appears to have no relationship with measures of urbanization and human presence.

**Figure S3.**
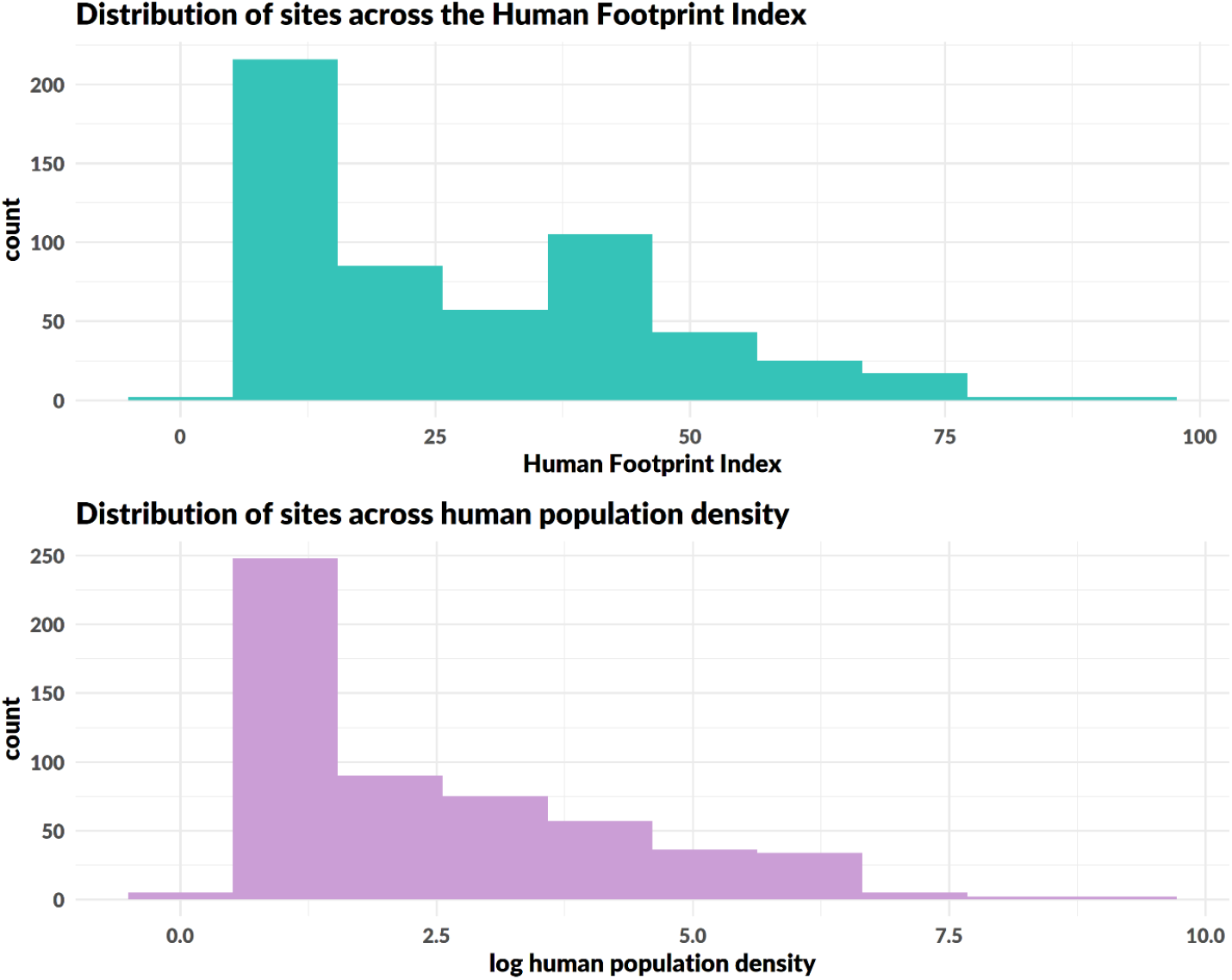
Distribution of sites across continuous measures of urbanization and human influence. A majority of sites are located in low to moderately transformed habitats.

**Table S1.**
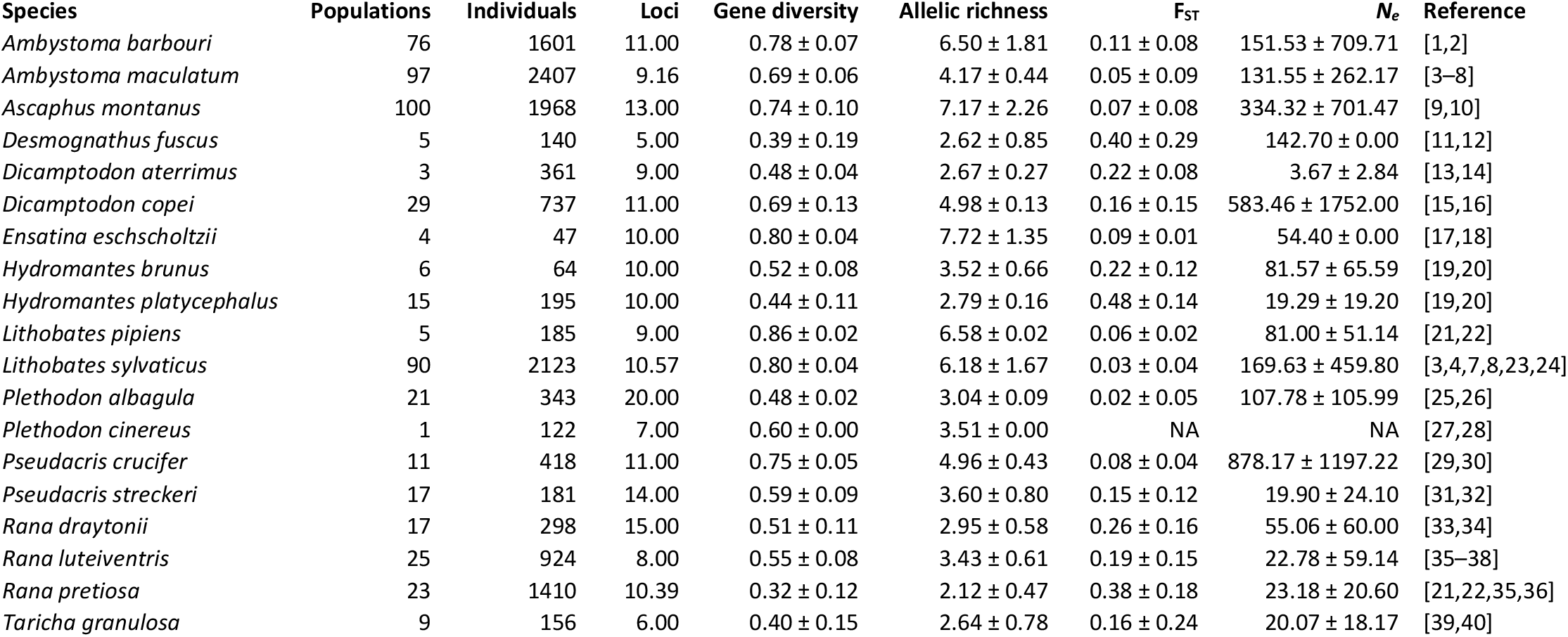
Data synthesis summary. Raw microsatellite data was obtained from 19 studies. **Populations:** the number of populations with >5 individuals included in analyses. **Individuals**: the number of individuals summed across all populations. **Loci:** average number of loci sampled across populations. Mean and standard deviations are presented for gene diversity, allelic richness, population-specific F_ST_, and effective population size (*N_e_*).

